# The 3'tsRNAs are aminoacylated: Implications for their biogenesis

**DOI:** 10.1101/2020.12.15.422816

**Authors:** Ziwei Liu, Hak kyun Kim, Jianpeng Xu, Mark A. Kay

## Abstract

Emerging evidence indicates that tRNA-derived small RNAs (tsRNAs) are involved in fine-tuning gene expression and become dysregulated in various cancers. We recently showed that the 22nt LeuCAG tsRNA from 3’ end of tRNA^Leu^ is required for efficient translation of a ribosomal protein mRNA and ribosome biogenesis. Inactivation of this tsRNA induced apoptosis in rapidly dividing cells and suppressed the growth of a patient derived orthotopic hepatocellular carcinoma in mice. The mechanism involved in the generation of the 3’-tsRNAs remains elusive and it is unclear if the 3’-ends of 3’-tsRNAs are aminoacylated. Here we report an enzymatic method utilizing exonuclease T to determine the 3’charging status of tRNAs and tsRNAs. Our results showed that the LeuCAG 3’-tsRNA is fully charged and originated solely from charged mature tRNA. When the leucyl-tRNA synthetase was knocked down, less tsRNA was generated while the mature tRNA was not reduced further supporting that tsRNA generation is regulated. The fact that the 3’-tsRNA is aminoacylated has implications for their biogenesis and provides additional insights into their biological role in post-transcriptional gene regulation.

## Introduction

Transfer RNAs (tRNAs) are cloverleaf molecules that bring amino acids to ribosomes for translating messenger RNA into protein. In addition to their roles in protein synthesis, recent years have seen emerging evidence of noncanonical regulatory functions of tRNA and tRNA fragments in regulatory events involved with maintaining homeostasis in varying cellular environments and various diseases including cancer, neurodegenerative disease, and viral infection (Balatti, Nigita et al., 2017, Blanco, Dietmann et al., 2014, Honda, Loher et al., 2015, Pekarsky, Balatti et al., 2016, Raina & Ibba, 2014, Ruggero, Guffanti et al., 2014). The tRNA fragments reported thus far can be classified into three groups: pre-tRNA fragments from the 5’ or 3’ portions of pre-tRNAs, tRNA halves that are cleaved in the anticodon loop of mature tRNAs, and tRNA-derived small RNAs (tsRNA) around 18-23nt from 5’ or 3’ of mature tRNAs (Gebetsberger & Polacek, 2013, Haussecker, Huang et al., 2010).

As key components of the translation system, tRNAs are highly regulated during the steps of maturation, modification, and trafficking (Kirchner & Ignatova, 2015). Under normal growth condition, high levels of aminoacylation of mature tRNAs are maintained through interactions with their cognate aminoacyl-tRNA synthetases (ARSs), which charge a specific tRNA with the correct amino acid, and translation elongation factors (Evans, Clark et al., 2017). While in mammalian cells, most of the mature tRNAs are complexed with ARSs, elongation or initiation factors and ribosomes, certain tRNAs are fragmented and released from the protein synthesis cycle (Schimmel, 2018). The accumulation of tRNA halves by angiogenin under stress conditions, interaction with Y-box-binding protein 1 (YBX1), and the inhibition of translation process by tRNA halves have been observed for years and studied extensively (Honda et al., 2015, Kumar, Kuscu et al., 2016, Lee & Collins, 2006, Yamasaki, Ivanov et al., 2009). In contrast, the biogenesis and regulation of tsRNAs, especially 3’-tsRNAs, remain largely elusive. The exploration of the tsRNA biogenesis pathways and the identification of the tsRNA interaction partners require further characterization of the tsRNAs.

Previously, we have shown that the 22nt 3’-tsRNA, tsRNA^Leu^-CAG, is required for efficient translation of several ribosomal protein mRNAs (Kim, Fuchs et al., 2017). Particularly, tsRNA^Leu^ is shown to facilitate the ribosomal protein RPS28 mRNA translation by pairing and altering the secondary structure of the mRNA. Northern blot results, sequencing and bioinformatic analysis showed that tsRNA^Leu^-CAG is a 22nt small RNA containing the universal CCA terminal sequence at its 3’ end. Unlike the tRNA halves, tsRNA^Leu^-CAG can be detected in mammalian cells under normal growth conditions, when most of the tRNA^Leu^ are charged. Thus, the charging status of tsRNA^Leu^ may offer a hint to their biogenesis and regulation. Nevertheless, several traditional methods used for analysis of tRNA are not feasible when applied to characterize the charging status of 3’-tsRNAs (Kohrer & Rajbhandary, 2008, Rizzino & Freundlich, 1975, Varshney, Lee et al., 1991). One reason is that the acid urea polyacrylamide gels that can separate charged tRNA from uncharged tRNA in size is not successfully used for detecting tsRNAs, as a result of their low abundance *in vivo* and the relatively lower sensitivity in this type of gel in a northern assay. Secondly, the periodate oxidation, which causes a 1-2nt shortening of the 3’ uncharged tRNA in urea acrylamide gels, leads to the inevitable degradation of a small amount of tRNA. While these methods are not problematic in a tRNA study, it is challenging to differentiate the degraded tRNA portion from the endogenous tsRNA.

Here, we report a new method employing the enzyme exonuclease T, which specifically cleaves the – A residue off the unprotected –CCA at the 3’ end of tsRNA, leaving the charged –CCA end intact by virtue of the acetylated protection of the 3’-aminoacyl group (Zuo & Deutscher, 2002). To begin to explore the implication of the tRNA charging status on tsRNA biogenesis, we studied the roles of the ARSs in regulating the formation of tsRNAs. We used various approaches to lower cytoplasmic leucyl-tRNA synthetase (LARS1) levels and/or activity in HeLa cells to alter the charging ratio of tRNA^Leu^ and evaluated their effects on tsRNA^Leu^ charging.

## Results

### Exonuclease T cleaves 3’ end adenosine from uncharged tRNA, whereas *N*-acetylated aminoacyl-tRNAs are protected

To test the charging status of a 3’-tsRNA^Leu^, exonuclease T was used under nearly neutral pH conditions to differentiate charged tsRNAs from uncharged tRNAs on a polyacrylamide urea gel. As shown in Fig. 1A, total RNAs extracted under acidic pH from HeLa cells were separated into two aliquots. The RNA in one aliquot was *N*-acetylated. The *N*-acetylation of the amino acid group attached to the tRNA by acetic anhydride stabilizes the aminoacyl bond and prevents the spontaneous deacylation reaction (Suzuki, Ueda et al., 1996) allowing protection of 3’ end at a higher pHs. The other aliquot was incubated under pH 9.0 as commonly used in the tRNA deacylation reaction. After the *N*-acetylation or the deacylation of the aminoacyl group, total RNA was treated with exonuclease T, which specifically digests single-stranded RNA from an unprotected 3’ end and stops at the -CC sites. Thus, uncharged tRNAs were one nucleotide shorter than the charged tRNAs after enzymatic digestion. Indeed, the tRNA^Leu^ band in the deacylated RNA sample was 1nt lower on an acrylamide gel than the tRNA^Leu^ band in the *N*-acetylation protected RNA sample as shown in Fig. 1B. Without the *N*-acetylation protection, the aminoacylated tRNAs were gradually deacylated during 3’ exonuclease digestion. As shown in Fig. 1C, *E. coli* leucyl-tRNA prepared by *E. coli* ARSs was deacylated within 30 mins and digested by exonuclease T thereafter, whereas the acetylated *E. coli* leucyl-tRNA remains intact after 60 mins. Taken together, this enzymatic method combining characteristics of exonuclease T and *N*-acetylation protection could be used to examine the 3’ status of tRNA or tRNA fragments containing a CCA end.

**Figure 1.**
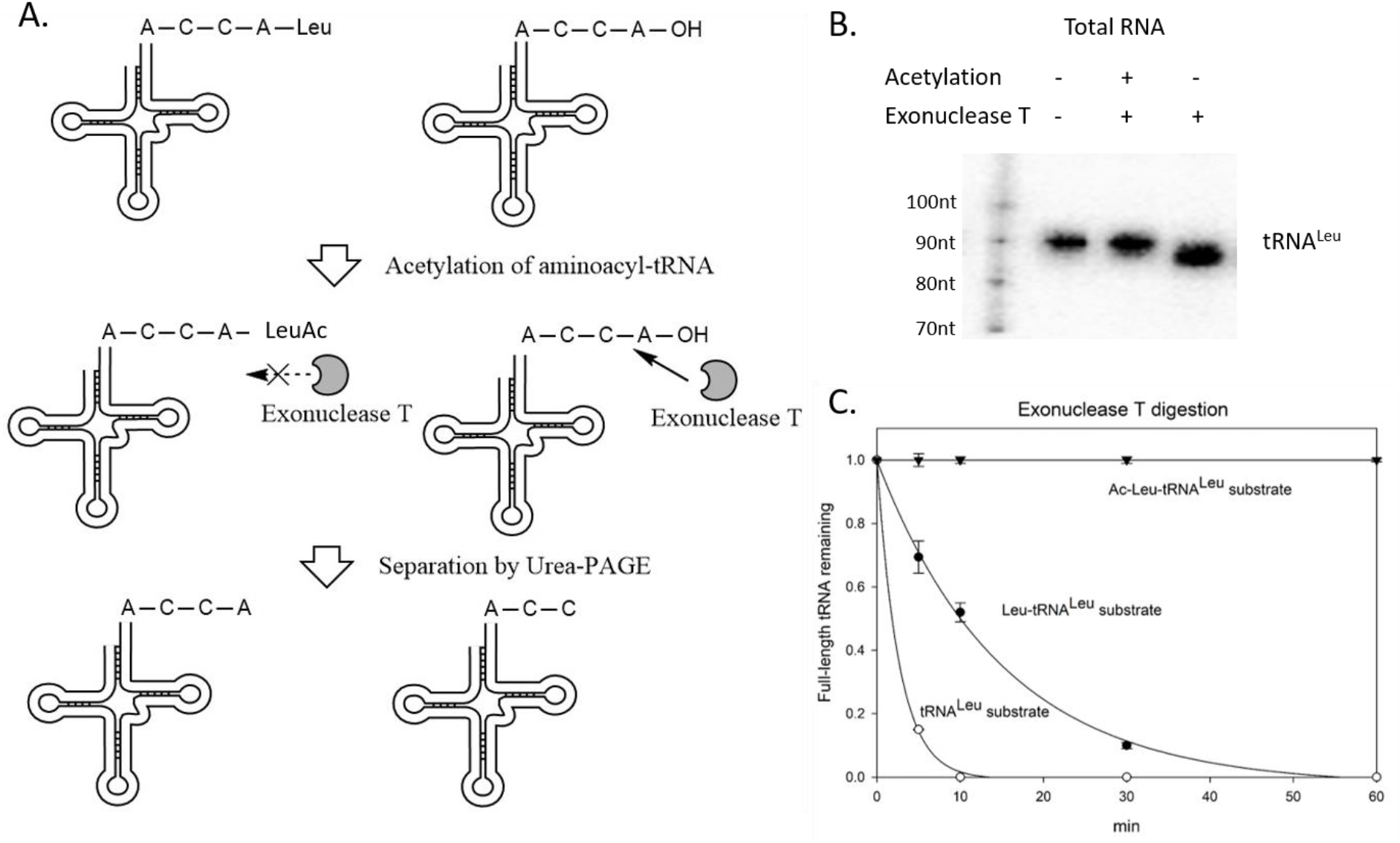
Exonuclease T digests the nucleotide A from 3'-end of uncharged tRNA. (A) Illustration of the enzymatic method to differentiate charged and uncharged tRNAs. (B) Uncharged tRNA^Leu^ digested by exonuclease T is 1 nt shorter than charged tRNA^Leu^ in a northern blot. Total RNA from HeLa cells were treated with or without acetylation, with or without exonuclease T digestion, and loaded on a 15% urea acrylamide gel. Radiolabeled RNA marker with various sizes were loaded in the left lane. A northern blot was performed using the tRNA^Leu^ probe. (C) Acetylation protects Leu-tRNA^Leu^ from deacylation and digestion with exonuclease T treatment. *E. coli* acetylated-Leu-tRNA^Leu^ (inverted triangle). Leu-tRNA^Leu^ (solid circle), uncharged tRNA^Leu^ (open circle) were prepared and subjected to exonuclease T digestion at room temperature. Samples were taken and quenched on ice at time points of 0, 5, 10, 30, 60 mins. Northern blots were used to detect the 1nt size difference after digestion. Levels of digested versus undigested tRNAs were quantified by Quantity One.

### TsRNA^Leu^ is fully charged regardless of the tRNA^Leu^ charging status

Utilizing our enzymatic method, we successfully characterized the charging status of this tsRNA. As shown in Fig. 2A, tsRNA^Leu^ was fully charged under normal cell growth condition, whereas the hsa-mir-223 miRNA control ending with -CCA, was digested by exonuclease T. To investigate whether the tRNA charging level affects the tsRNA charging level, HeLa cells were treated with the leucyl-tRNA synthetase aminoacylation inhibitor, leucinol. Leucinol is a leucine mimic (Fig. 2B) that binds to the active site of LARS1 and competes with leucine for aminoacylation (Bonfils, Jaquenoud et al., 2012, Vaughan & Hansen, 1973). Adding leucinol to HeLa cell culture resulted in a decrease of tRNA^Leu^ charging to about fifty percent, while the tsRNA^Leu^ was still fully charged under this condition (Fig. 2C). Accordingly, the fact that tsRNAs are fully charged and not related with tRNA charging levels indicates that the tsRNA^Leu^ is generated from the mature tRNA after mature tRNA is charged by the aminoacyl-tRNA synthetases.

**Figure 2.**
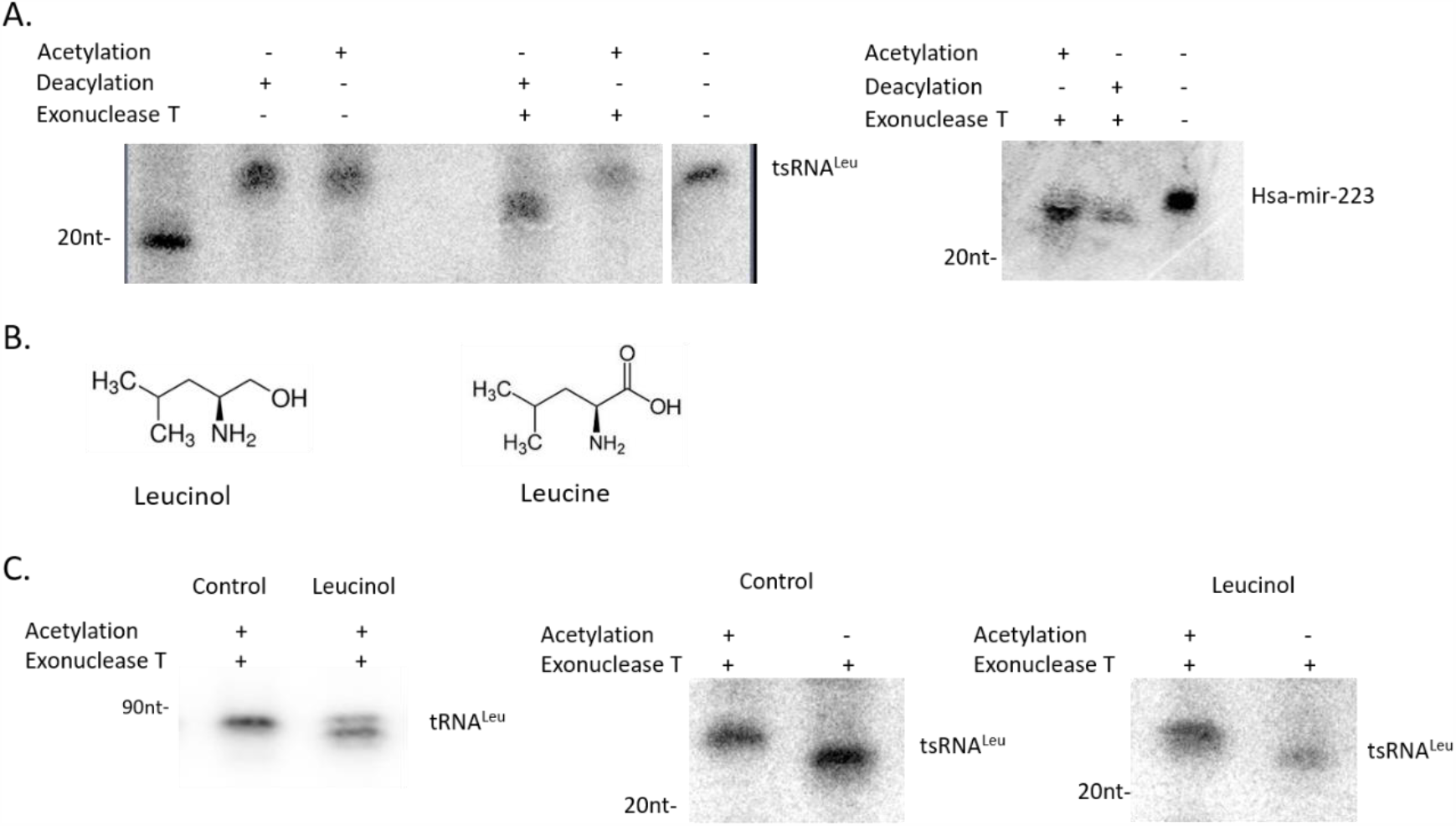
TsRNA^Leu^ are fully charged under various growth conditions. (A) Under normal growth condition, tsRNA^Leu^ are aminoacylated. HeLa total RNA extracted under pH 5.0 were either acetylated or deacylated, ethanol precipitated, and exonuclease T digested. Size marker was loaded on the left lane and total RNA without any treatment were used for comparison on the right lane. DNA probes antisense to tsRNA^Leu^ were used for northern blot (left panel). The hsa-miR-223-3p miRCURY LNA miRNA (Qiagen) detection probe was used for detection of this miRNA ending with CCA in sequence (right panel). (B) Leucinol is an analog of leucine and inhibitor of LARS1. (C) tsRNA^Leu^ are fully charged under leucinol condition. Total RNA of HeLa cells grown with or without leucinol (2 mM) were extracted and examined for charging status. Full-length tRNA^Leu^ were half charged under this condition (left) while tsRNA^Leu^ is fully charged (middle and right).

### The tsRNA^Leu^ concentration is regulated by LARS1

To explore the effect of leucyl-tRNA synthetase (LARS) on tsRNA formation, we used siRNAs to separately reduce the cytoplasmic LARS1 and the mitochondrial LARS2, and then determined the tsRNA levels in each case. As shown in Fig. 3A, knocking down of LARS1, but not LARS2, led to a reduction in the tsRNA level. In order to examine whether or not the decrease in the tsRNA level was associated with a lowering in the tRNA charging level, we measured the charging levels of tRNA and tsRNA in the cells transfected with siRNAs directed against LARS1. Northern blot results showed that the tRNA^Leu^ was still fully charged with the remaining LARS1, and the tsRNA^Leu^ was also fully charged under this condition (Fig. 3B). These results suggest that the decrease in tsRNA level was not the result of a limited Leu-tRNA^Leu^ and supports the idea that the LARS1 is part of the tsRNA^Leu^ biogenesis pathway. The fact that the tsRNA^Leu^ level is regulated through LARS1, but not LARS2, correlates with the exclusive cytoplasmic localization of 3’ tsRNAs (Haussecker et al., 2010).

**Figure 3.**
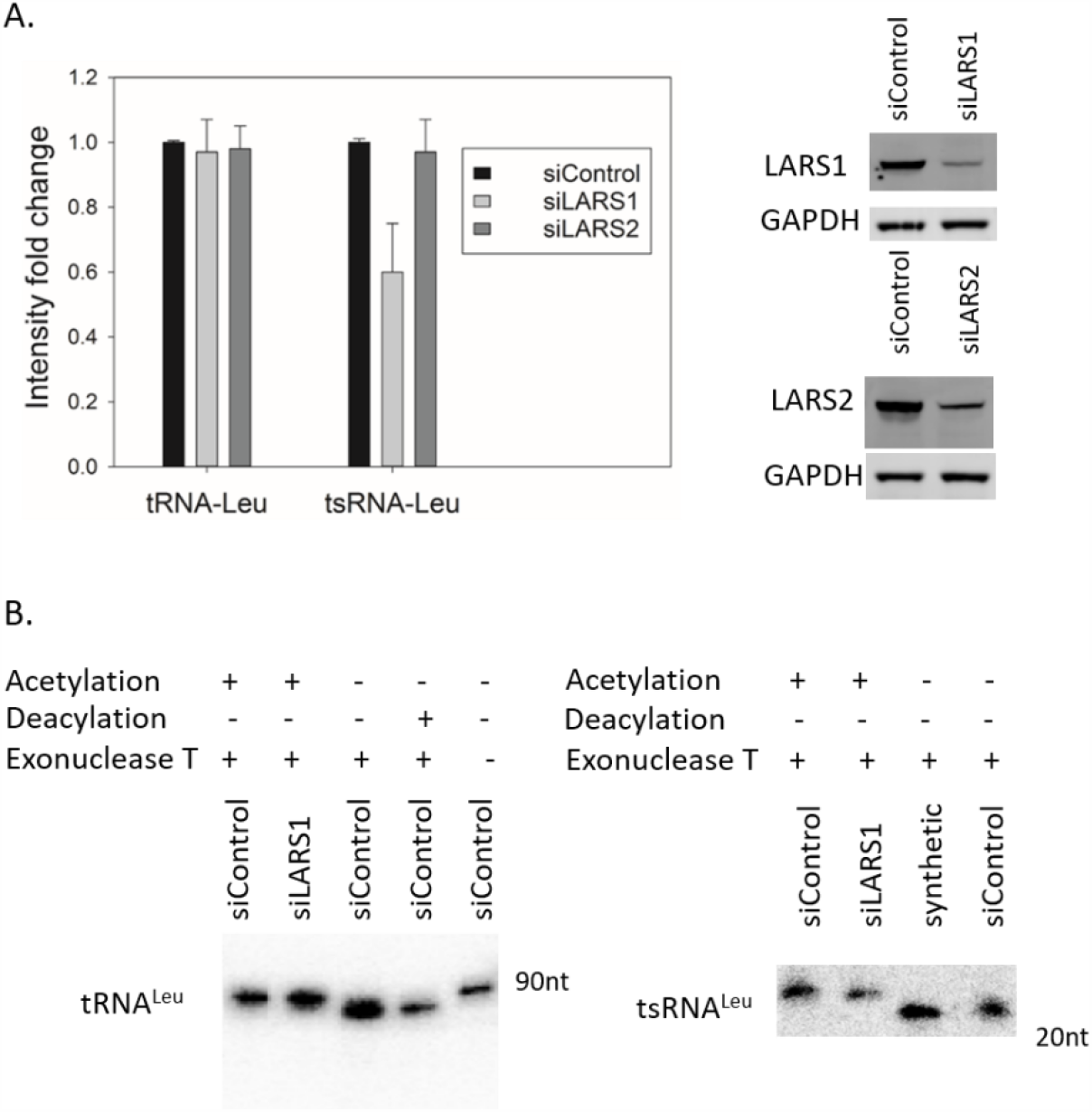
Knocking down of LARS1 decreases tsRNA^Leu^. (A) SiRNAs targeting either LARS1 or LARS2 were transfected by Lipofectamine 3000 to HeLa cells in a 6-well plate. Total RNAs were extracted after 72 hours, and northern blots were performed to quantify the tRNA^Leu^ and tsRNA^Leu^ levels. Experiments were conducted triple times independently. Western blots were used to test full-length LARS1 and LARS2 expression levels. (B) tRNA^Leu^ and tsRNA^Leu^ are both fully charged in cells transfected either with LARS1 siRNA (siLARS1) or negative control siRNA (siControl). A chemically synthetic 22-mer tsRNA oligo was treated and used as a size control.

### Overexpressing tRNA^Leu^ charging variants give rise to fully charged tsRNA^Leu^

We wanted to determine if impaired tRNA charging could affect the generation of tsRNA, and hence constructed various plasmids that express the wild-type and variant tRNAs (Fig. 4A). Overexpression of wild-type tRNA^Leu^ resulted in an increase in the tsRNA^Leu^ (Fig. 4B). We constructed tRNA^Leu^ overexpression plasmids with various mutations in the tRNA identity elements known to affect tRNA charging (Fig. 4A). After each plasmid was transfected separately, we determined the tsRNA charging levels. Human tRNA^Leu^ A73, as reported previously, was known to significantly affect the LARS1 recognition. Among the human tRNA^Leu^ A73 mutations, the A73C mutant cannot be charged by LARS1 *in vitro*, and the uncharged tRNA^Leu^ A73C mutant also inhibits the aminoacylation of wild-type tRNA^Leu^ (Breitschopf & Gross, 1994). As shown in Fig. 4B, overexpression of tRNA^Leu^ A73C results in an increased amount of uncharged tRNA^Leu^ yet there was no change in the tsRNA charging level.

**Figure 4.**
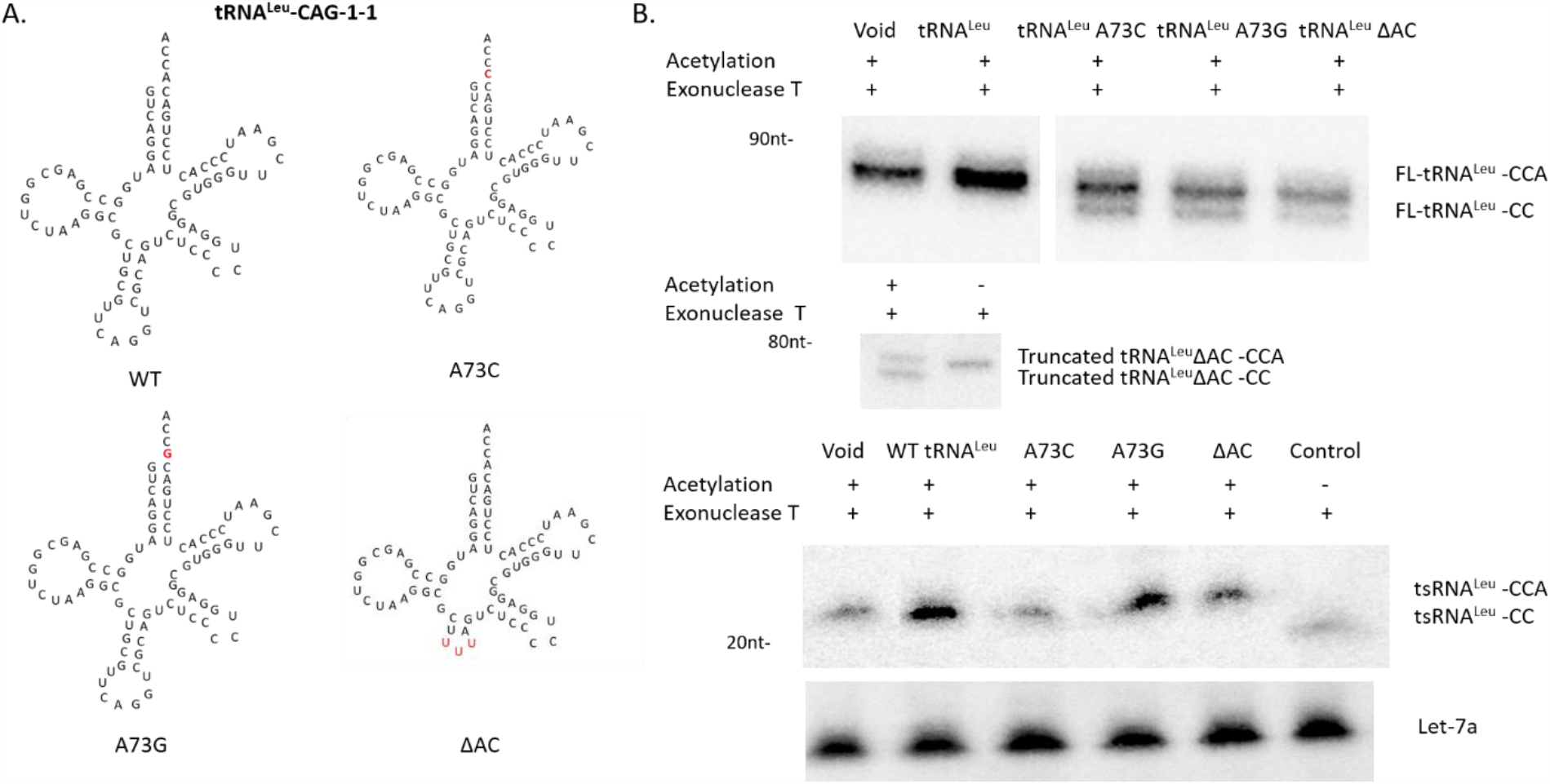
Overexpression of chargeable or unchargeable tRNA^Leu^ give rise to fully charged tsRNA^Leu^. (A) Sequence and secondary structure of human tRNA^Leu^ and variants. (B) Charging levels of tRNA^Leu^ and tsRNA^Leu^ in the HeLa cells transfected with plasmids carrying various tRNA^Leu^ mutation. ΔAC stands for the tRNA^Leu^ truncated variant lacking the anticodon domain. Overexpression of ΔAC variant leads to extra truncated tRNA bands around 70nt size in northern blot as shown in the middle panel. Let-7a-3p was used as loading control.

Another mutant tRNA^Leu^ A73G was previously shown to confer serine acceptance by seryl-tRNA synthetase (SerRS) instead of LARS1 *in vitro* (Breitschopf, Achsel et al., 1995). Transfection of tRNA^Leu^ A73G resulted in an increased level of charged tRNA^Leu^ and a small fraction of uncharged tRNA^Leu^, possibly due to the impaired mis-aminoacylation efficiency by SerRS. In this case, the tsRNA^Leu^ remained fully charged with a slightly elevated tsRNA concentration level compared to the A73C mutant. This suggests that the tsRNA was also generated from the mischarged tRNA. To clearly differentiate the overexpressed tRNA from the endogenous tRNA, a plasmid encoding truncated mutant of tRNA^Leu^ lacking the anticodon loop was designed and transfected. Here we showed that the ΔAC tRNA^Leu^ was not efficiently charged *in vivo* (Fig. 4B), despite the previous implications that the tRNA^Leu^ anticodon loop is dispensable for the human LARS1 aminoacylation (Breitschopf et al., 1995). This impaired aminoacylation phenomenon is similar to the anticodon loop deletion mutant in *E. coli* (Larkin, Williams et al., 2002). As a result, it seems there are differences in the hLARS1 charging activity between living cells and a cell-free system. As shown in Fig. 4B, the tsRNA^Leu^ was still fully charged with tRNA^Leu^ ΔAC overexpression. Taken together, all of these studies are supportive of the formation of tsRNA^Leu^ requires a charged mature tRNA.

### Bioinformatic analysis of tsRNA^Leu^ in small RNA sequencing libraries

To evaluate the tsRNA level from high-throughput sequencing databases, the phosphorylated status of the 5’ end of tsRNA^Leu^ was examined by several enzymatic methods. As shown in Fig. 5A., total RNA was treated with either T4 polynucleotide kinase (T4 PNK) or fast alkaline phosphatase (FAP), and the mobility of tsRNA^Leu^ bands was tested by northern blot. The FAP treatment was able to remove the 5’ phosphate group of tsRNA and shift the band, whereas the T4 PNK treatment that phosphorylates the 5’ end of RNA did not affect the band position. In addition, the terminal exonuclease that is able to digest only 5’ phosphorylated RNAs removed the tsRNA^Leu^ band as shown in Fig. 5B. Similar results were observed using a let-7a miRNA probe. Taken together, these results suggest that the 5’ end of tsRNA^Leu^ is phosphorylated and would be amenable to 5’ adaptor ligation used in a high-throughput sequencing.

**Figure 5.**
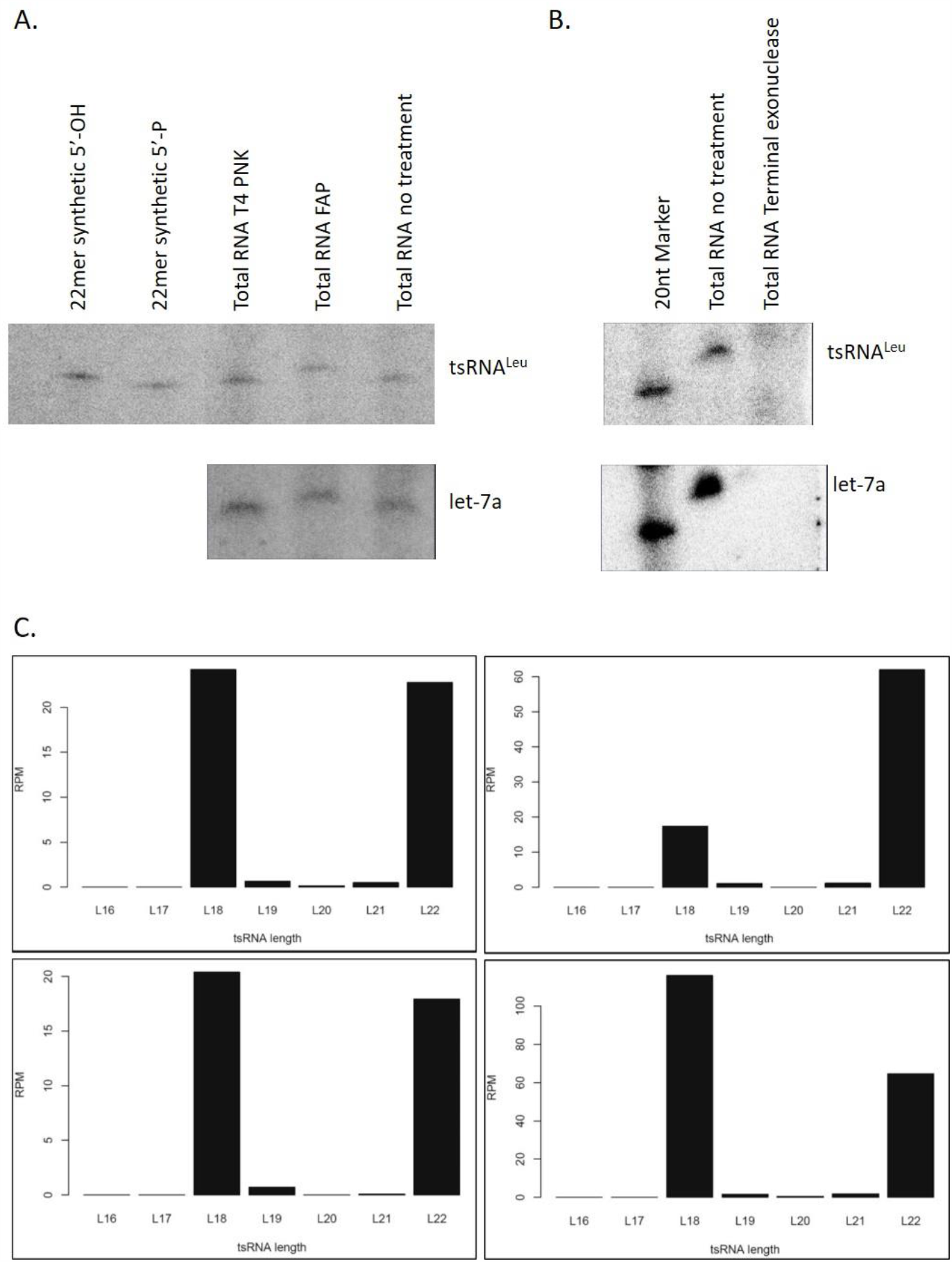
5' terminal 3' tsRNA^Leu^ is phosphorylated. (A) Hela total RNA was treated with T4 PNK or FAP and northern blots were performed. Nontreated RNA sample, synthetic 22mer tsRNA^Leu^ with either a 5'-hydroxyl or 5'-phosphate group were used as control to indicate the upshift of tsRNA^Leu^ after FAP treatment. MiRNA let-7a serves as a control RNA with 5'-phosphate. (B) Terminal exonuclease digested the tsRNA^Leu^ as well as let-7a with the 5'-phosphate. (C) Length distribution of reads that mapped to 3' tRNA^Leu^-CAG-1-1. TsRNA^Leu^ reads from Hela small RNA libraries GSM876014, GSM876013, GSM876017, GSM876016.

In most RNA sequencing libraries, tRNA reads may not be accurately quantified because the reverse transcriptase tends to stop at various modification sites and/or by structural elements, resulting in truncated tRNA reads. To exclude the truncated reads from full-length tRNAs, only small RNA sequencing libraries with size selection between 17-30 bases were explored for our study. We analyzed four different small RNA libraries from HeLa cells and aligned the reads to the tRNA^Leu^-CAG-1-1 gene according to the hg19 genomic tRNA database (Chan & Lowe, 2009). As shown in Fig. 5C, results from all sequencing databases showed two specific peaks with a similar number of reads in each sample, representing the 22nt full-length tsRNA^Leu^ and the 18nt truncated tsRNA, the latter, which was not detected by northern blot (Kim et al., 2017). Although most truncated reads in deep sequencing are thought to be from reverse transcriptase stops at the modification site, we cannot rule out the possibility that there are different levels of degradation of tsRNA at the 18nt modification (m1A) site during the adaptor ligation steps. In either case, the sum of the two peaks can be calculated to determine the relative tsRNA^Leu^ concentration. In addition, the construction of small RNA libraries in the past seldom involves the tRNA deacylation step. It is possible that the 3’ end of the tRNAs and tsRNAs would not be ligated to the adaptor due to the presence of the aminoacyl bonds at the time of library preparation. In summary, determining the correlation of the 22-nt tsRNA with cellular status by high throughput sequencing method requires further evaluating tRNA modification levels and developing optimized tRNA-seq protocols including removing modifications and eliminating the ligation (Cozen, Quartley et al., 2015, Gogakos, Brown et al., 2017, Hrabeta-Robinson, Marcus et al., 2017, Zheng, Qin et al., 2015).

## Discussion

As a novel class of small ncRNAs discovered in recent years, the reporting functions of the tsRNAs in translational control, oncogenesis, and neurologic disorders are emerging (Schimmel, 2018). Dysregulation of tsRNAs have been found in various cancers and diseases (Goodarzi, Liu et al., 2015, Pekarsky et al., 2016) and in order to gain some insight into how, where and when they are generated, we sought to determine if the 3’-tsRNAs are aminoacylated. Because of the low abundance of 3’-tsRNA and instability of the phosphodiester bond in neutral or high pH solutions and elevated temperature in beta elimination treatment, characterizing the charging status of 3’-tsRNA^Leu^ has been difficult. Here we developed a sensitive enzymatic method to examine the charging status of 3’-tsRNA^Leu^ under various cellular conditions. Elucidating the fact that 3’-tsRNA^Leu^ are fully charged albeit of tRNA charging level not only suggests that tsRNA^Leu^ is generated from mature aminoacyl-tRNA, but also indicates the interaction of tsRNA with ARSs (Fig. 6). In total, our study characterizing the terminal status of tsRNA, together with manipulation of aminoacyl-tRNA synthetases, provide new insights into the generation of 3’-tsRNAs.

**Figure 6.**
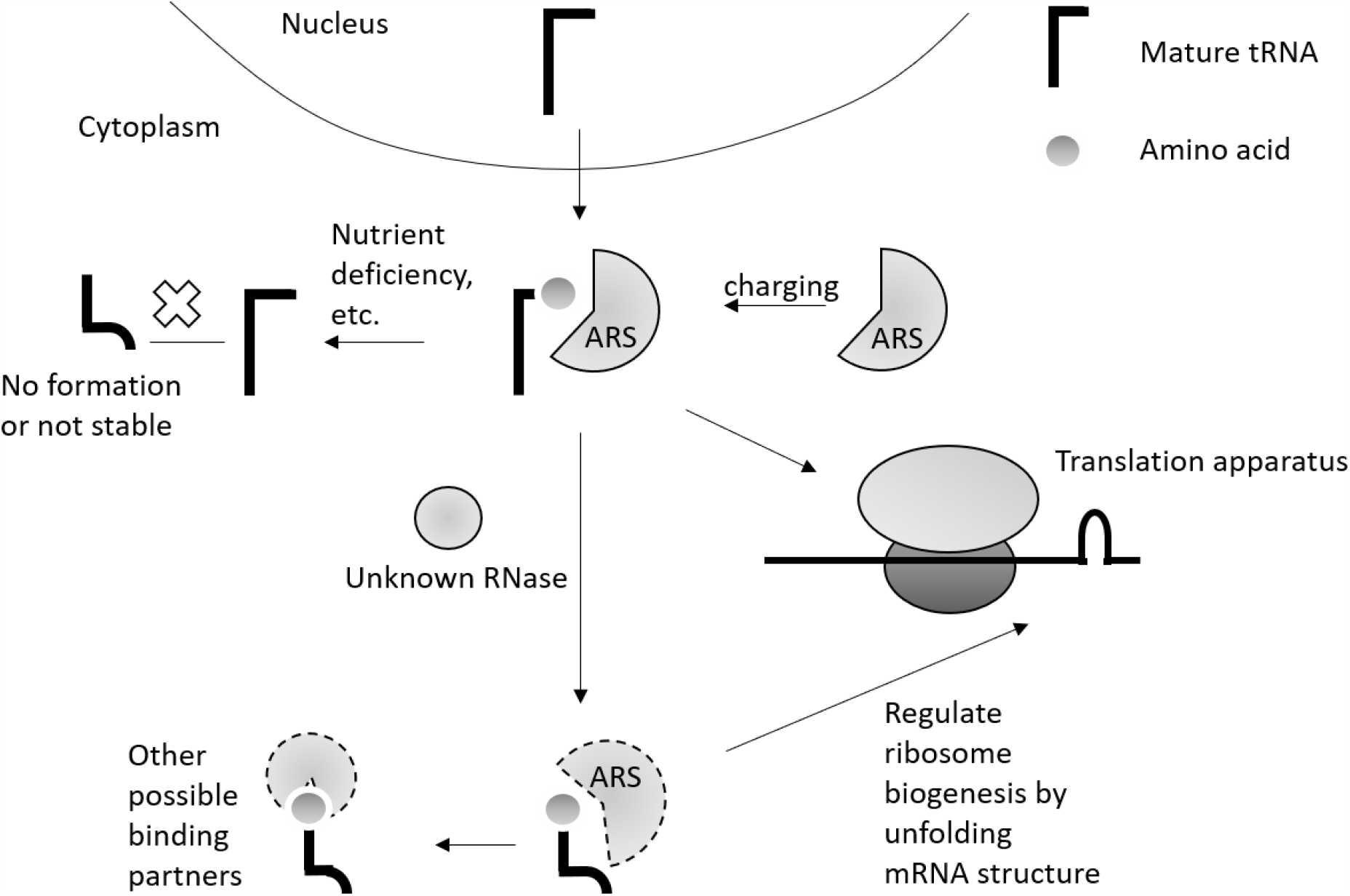
3'-tsRNAs were generated from cytoplasmic aminoacyl-tRNAs and regulated by aminoacyl-tRNA synthetases (ARS).

Since it has been shown that tsRNAs from 3’ end of tRNAs are localized in the cytoplasm (Haussecker et al., 2010), it is not surprising that tsRNA^Leu^ is charged under normal growth conditions when most tRNAs are fully aminoacylated. Nonetheless, lowering the aminoacylation level of tRNA^Leu^ by leucinol to fifty percent still gives rise to fully charged tsRNAs. This result indicated that tsRNA^Leu^ originates from mature tRNAs and the processing of tsRNA^Leu^ happens after aminoacylation by ARSs. Further evidence came from *in vivo* overexpression of wildtype tRNA^Leu^ and various mutants. While overexpression of wild-type tRNA^Leu^ CAG-1-1 gene elevates the level of charged tsRNA^Leu^ compared to the control, overexpression of the unchargeable mutant, A73C leads to no change or a slightly lower amount of aminoacyl-tsRNA. The effect of A73C overexpression results from the inhibition of wild-type tRNA^Leu^ aminoacylation induced by the presence of A73C tRNA^Leu^. In comparison, overexpression of the chargeable mutant A73G tRNA^Leu^ led to the overexpression of tsRNA as in the case with the wild-type tRNA gene, but to a lower extent. A73G tRNA^Leu^ was shown to be mischarged by SerRS *in vitro*, suggesting the biogenesis of this tsRNA is not limited by the fidelity of tRNA aminoacylation. In addition, by deleting the anticodon loop of tRNA^Leu^, we established that the ΔAC tRNA^Leu^ is charged to fifty percent by LARS1 *in vivo* and gave rise to increased tsRNA concentrations. Taken together, these data strongly suggest that 3’-tsRNA^Leu^ is generated only from aminoacylated tRNAs, and that elevated chargeable tRNAs might regulate the production of more 3’-tsRNA^Leu^.

tRNA charging occurs via ARS proteins in a poorly defined complex referred to as the multi-aminoacyl-tRNA synthetase complex (MSC) consisting of multiple proteins including elongation factor 1A. This along with a number of studies have shown that the MSC is associated with actively translating ribosomes (Kong & Kim, 2020). Reducing LARS1 did not result in an increased uncharged tRNA^Leu^ but rather resulted in a decrease in the tsRNA^Leu^ level, suggesting that the tsRNA^Leu^ biogenesis could in fact be regulated by aminoacyl-tRNA synthetases. Evidence of tsRNA binding with ARSs further supported the possibility that at least a portion of the tsRNAs is contained near the translation apparatus (Mleczko, Celichowski et al., 2018). The finding that tRNA fragments are associated with ARSs, but not EF1A further supports the idea that tRNA derived small RNAs might be regulated by the ARSs level (Saikia, Krokowski et al., 2012).

Because the 3’-tsRNA^Leu^ regulates ribosome biogenesis by regulating the translation of a key ribosomal protein RPS28 (Kim et al., 2017, Kim, Xu et al., 2019), these complexes on actively translating ribosomes may indeed function in some sort of complex feedback regulatory mechanism. It is possible that the responsible ribonucleases for tsRNA formation interact with the aminoacyl-tRNA synthetases or ribosomes to regulate the ribosome biogenesis and other cellular processes as part of the feedback mechanism.

It has been reported that tRNA fragments could be generated by various RNases, such as RNase T2, angiogenin (ANG), and Dicer (Cole, Sobala et al., 2009, Kumar, Anaya et al., 2014). Nevertheless, these reported enzymes are responsible for the formation of only certain species of the tRNA fragments. Specifically, the biogenesis of 22mer 3’-tsRNAs from mature tRNAs are not well determined. Firstly, human RNase T2 was shown to digest ssRNA between purine and uridine residues, leaving the RNA fragments ending with 2’, 3’-cyclophosphate configurations (Greulich, Wagner et al., 2019). The fact that the 3’-tsRNAs are cleaved between T54 and Ψ55 and their 5’-ends are phosphorylated suggests a mechanism other than RNase T2. Secondly, a previous enzymatic study had shown that ANG could generate tRNA fragments but differed in sizes from the 22-mer tsRNAs (Li, Ender et al., 2012). Recently a knock-out study also supported that angiogenin was responsible for certain species of tRNA halves, but not for the tsRNAs (Su, Kuscu et al., 2019). Thirdly, as a double-strand endonuclease, Dicer was shown to cut tRNA mainly at double-stranded T-arm region and minimally at the T-loop (Reinsborough, Ipas et al., 2019). Indeed, though the level of several 3’-tRNA fragments were shown to be Dicer-dependent, there were no differences in the level of other tsRNA species, such as tRNA^Leu^-CAG fragments, in knock-out cells (Li et al., 2012). Taken together, defining the cleavage mechanism of tsRNAs requires further characterization of alternate RNases, and it could be possible that multiple steps and enzyme pathways are involved in the biogenesis of tsRNA.

Changes in tRNA post-transcriptional modification level are reported to affect the biogenesis of tRNA fragments. It has been reported that lack of cytosine-5 methylation from DNMT2 or decreasing m1A and m3C levels by ALKBH3 could promote the production of tRNA halves by angiogenin (Blanco et al., 2014, Chen, Qi et al., 2019). In addition, the deficiency of TRMT10A, a guanosine 9 tRNA methyltransferase, led to the fragmentation of tRNA^Gln^ and tRNA^iMeth^ in pancreatic β-like cells (Cosentino, Toivonen et al., 2018). Recently, an RNA phospho-methyltransferase BCDIN3D is reported to interact with the tRNA^His^ 3’ fragments and regulate their biogenesis (Reinsborough et al., 2019). It is possible that the RNA modifying enzymes regulate the tsRNA generation by affecting the stability of the tRNA structure and the interaction among the tRNA, aminoacyl-tRNA synthetase complex and the ribosomes. Although we did not observe changes in the tsRNA level when knocking down the pseudouridine synthase TruB1, which is responsible for the Ψ55 modification (data not shown) (Gutgsell, Englund et al., 2000), we cannot rule out the effect of pseudouridine and other tRNA modification levels on tsRNA formation. Further identification of tRNA and tsRNA modification proteins may provide more evidence for the high-variable, cell-specific biogenesis of tsRNA.

## Materials AND Methods

### Cell culture and transfection

HeLa cells were cultured in Dulbecco’s modified Eagle’s medium supplemented with 10% fetal bovine serum in a 5% CO_2_ incubator. For RNA interference, Opti-MEM medium and Lipofectamine 3000 (ThermoFisher) was used as protocol. HeLa cells were transfected with control siRNA (non-targeting pool, Dharmacon) or siRNAs targeting human leucyl-tRNA synthetase 1 or 2 (Dharmacon) separately. For transfection of tRNA plasmid, HeLa cells were transfected by Lipofectamine 3000 with 2 μg of plasmid encoding tRNA wild-type or variant genes and incubated for 24 hours before RNA extraction.

### Construction of tDNA plasmids

The gene encoding human cytosolic tRNA^Leu^-CAG-1-1 was cloned with 150 bp 5’ and 3’ flanking regions from the genome. The corresponding tRNA^Leu^ region was inserted into pUC19 plasmid (ptRNA^Leu^CAG). Mutation of A73 to G or C, and the anticodon stem deletion in the tDNA genes was made by QuickChange lightning site-directed mutagenesis kit (Agilent) using the following primers:

CAG-AtoG_F ATCCCACTCCTGACGATATGTGTTTTCCGT

CAG-AtoG_R ACGGAAAACACATATCGTCAGGAGTGGGAT

CAG-AtoC_F ATCCCACTCCTGACCATATGTGTTTTCCGT

CAG-AtoC_R ACGGAAAACACATATGGTCAGGAGTGGGAT

CAG-delta_AC_F GGTCGCAGTCTTTTGGCGTGGGTTC

CAG-delta_AC_R GAACCCACGCCAAAAGACTGCGACC

### RNA isolation, N-acetyl protection and deacylation of tRNA, and exonuclease T treatment

Cells were harvested using trypsin and washed with cold phosphate-buffered saline. Total RNA was extracted by Trizol (Invitrogen) and Direct-zol RNA kit (Zymo Research). RNA was kept in buffers of pH 5 at all time and avoid too many freeze-thaw cycles to keep the aminoacyl group attached to tRNA. For aminoacylation characterizing assay, total RNA was separated into two fractions. One fraction was incubated on ice in 250 µl of 0.2 M sodium acetate with 4 µl acidic anhydride for 2 hours for *N*-acetylation as described previously (Walker & Fredrick, 2008). The other fraction was incubated at 37 degrees at pH 9.0 for 30 min to deacylated the aminoacyl group (Walker & Fredrick, 2008). Both fractions were then ethanol precipitated at -80 degrees for 2 hours and washed with 80% ethanol. 10 ug total RNA was resuspended in 87 µl RT-PCR grade water (ThermoFisher), and 3 µl exonuclease T with 10 µl buffer 4 (NEB) was used in the enzymatic reaction. The digestion reactions were kept at room temperature for 1 hour, and then the RNA clean and concentrator kit (Zymo) was used to remove the exonuclease T and other residues. The RNA samples were quantified by nanodrop (Agilent) and subjected to subsequent applications.

### Northern blot

10 µg total RNA samples were mixed with gel loading buffer II (ThermoFisher) and separated by 10% or 15% TBE-urea gels (Novex) in 1x Tris-borate-EDTA (TBE) buffer at 180 V for 90 min. RNA was transferred to a positively charged nylon membrane (Hybond N+, GE Healthcare) through semi-dry electrophoretic transfer cell (Biorad) in 0.5x TBE buffer at 5 Watt for 1 hour. UV crosslinker (Stratagene) was used to crosslink RNA to the membranes at 150 mJ cm^-2^. Membranes were prehybridized by PerfectHyb plus (Sigma) hybridization buffer for 1 hour and hybridized overnight with the [γ-32P] ATP-labeled probe which is complementary to 3’ end of tsRNA^Leu^-CAG-1-1. Membranes were washed with 2x SSC and 0.1% SDS three times for 10 minutes.

### Aminoacyl-tRNA preparation

Aminoacylation was performed in 100 mM Hepes-NaOH (pH 7.6), 30 mM KCl, 10 mM MgCl2, 1 mM dithiothreitol (DTT), 2 mM adenosine triphosphate (ATP), 10 mM Leucine, 1.5 mg/mL of *Escherichia coli* total tRNA (Roche), and 10 nM of crude ARSs from *Escherichia coli* (Sigma, A3646) (Splan, Musier-Forsyth et al., 2008, Walker & Fredrick, 2008). After a 30 min incubation at 37 °C, tRNA was ethanol precipitated at -80 degrees for 2 hours and washed twice with 70% ethanol. The resulted mixture of aa-tRNA and tRNA was separated and analyzed by urea acrylamide gels and northern blots.

### Western Blot

Transfected cells were lysed in cell lysis buffer (Cell signaling) with 1x protease inhibitor cocktail (Sigma) on ice for 15 minutes. After centrifugation at 13,000 rpm at 4 degrees for 30 minutes, precipitant was removed, and protein was quantified by Pierce BCA assay kit (ThermoFisher). 4 µg protein was resolved on NuPAGE 4-12% Bis-Tris protein gels (ThermoFisher) and transferred to PVDF membrane (Fisher). The membranes were blocked by Odyssey blocking buffer (Licor) and then immunoblotted with the cytoplasmic LARS1, the mitochondrial LARS2 or GAPDH primary antibodies (Sigma) and IRDye secondary antibodies (Licor). Results were visualized using the Odyssey imaging systems.

### 5' phosphorylation characterization

Total RNA was extracted by Trizol and treated with FastAP (ThermoFisher), T4 polynucleotide kinase (New England Biolabs, with 1mM ATP) and Terminal exonuclease (buffer A) following standard protocol. 5' hydroxyl and 5' phosphorylated 22mer tsRNA^Leu^ oligos were synthesized by Dharmacon.

### Bioinformatic analysis of 3'-tsRNA

The small RNA deep sequencing datasets for HeLa cell line were downloaded from the GEO database. Only libraries generated from 17-30 bases sized selected small RNAs were analysed. Human tRNA gene sequences (human hg19) were downloaded from the genomic tRNA database (27). Small RNA reads were mapped onto the tRNA gene sequences with CCA end added at the 3’ end using Bowtie version 1.2.2 (Langmead, Trapnell et al., 2009).

## Acknowledgements

This work was supported by NIH DK114483 (MAK).

## Author contributions

ZL, and MAK conceived and designed the experiments; ZL performed and analyzed experiments. HK provided reagents and resources. JP analyzed and interpreted the bioinformatic data. MAK supervised experiments and provided funding. ZL and MAK wrote the manuscript with input from all other authors.

